# X-chromosome loss rescues Sertoli cell maturation and spermatogenesis in Klinefelter syndrome

**DOI:** 10.1101/2024.02.09.576184

**Authors:** Sofia B. Winge, Niels E. Skakkebaek, Lise Aksglaede, Gülizar Saritaş, Ewa Rajpert-De Meyts, Ellen Goossens, Anders Juul, Kristian Almstrup

## Abstract

Klinefelter syndrome (47,XXY) causes infertility with a testicular histology comprising two types of Sertoli cell-only tubules, representing mature and immature-like Sertoli cells, and occasionally focal spermatogenesis. Here, we show that the immature Sertoli cells highly expressed *XIST* and have two X-chromosomes, while the mature Sertoli cells lack *XIST* expression and have only one X-chromosome. Sertoli cells supporting focal spermatogenesis also lack *XIST* expression and the additional X-chromosome, while the spermatogonia expressed *XIST* despite having only one X-chromosome. *XIST* was expressed in Sertoli cells until puberty, where a gradual loss was observed. Our results suggest that a micro-mosaic loss of the additional X-chromosome is needed for Sertoli cells to mature and to allow focal spermatogenesis.

## Introduction

Klinefelter syndrome (KS) is caused by a 47,XXY karyotype (Jacobs and Strong, 1959) and has an estimated prevalence of 152 per 100,000 new-born boys (Gravholt et al., 2018) but with only around 25-40% ever receiving the diagnosis (Berglund et al., 2019; Bojesen et al., 2003; Herlihy et al., 2011; Zhao et al., 2022). The phenotype varies considerably with the most persistent symptoms being small testes, non-obstructive azoospermia, and hypergonadotropic hypogonadism (Zitzmann et al., 2021). In addition, men with KS are at increased risk of developing osteoporosis, obesity, type II diabetes and cardiovascular disease as well as cognitive and psycho-social difficulties (Bojesen and Gravholt, 2011; Ross et al., 2012). The testicular histology of men with KS comprises Leydig cell hyperplasia, hyalinized “ghost” tubules, and tubules containing only Sertoli cells (Aksglaede et al., 2006; Winge et al., 2020). Occasionally, focal spermatogenesis can be observed, giving the patient possibility to father a biological child, usually with the help of testicular sperm extraction (TESE) and subsequently intracytoplasmic sperm injection (ICSI) (Corona et al., 2017). It remains unknown what biologically determines if focal spermatogenesis can occur.

In 46,XX females, one of the two X-chromosomes undergoes inactivation to balance the dosage of X-linked genes, a process orchestrated by the long non-coding RNA, *XIST* (Galupa and Heard, 2015; Lee, 2011). An inactivated X-chromosome can be seen in somatic cells as a dense Barr body (Barr and Bertram, 1949; Ohno et al., 1959). The inactivation is not complete, though, as around 15% of X-chromosomal genes escape inactivation (Berletch et al., 2011). Since *XIST* is expressed at high levels in men with KS (reviewed in Winge et al., 2020), the additional X-chromosome is expected to undergo inactivation as well.

With the emergence of single-cell RNA sequencing (scRNAseq), three studies including samples from a total of six men with KS (Laurentino et al., 2019; Mahyari et al., 2021; Zhao et al., 2020), all indicated a central role of the Sertoli cells in the testicular pathology of men with KS (Winge et al., 2020), which supports earlier morphological studies from 1969 (Skakkebaek, 1969). In the latter, two distinct types of Sertoli cell-only (SCO) tubules, termed type A and B, were identified, with type A Sertoli cells resembling adult mature Sertoli cells and type B Sertoli cells resembling immature Sertoli cells (Skakkebaek, 1969). Whereas Barr bodies were never observed in type A tubules, a Barr body was present in 16% of Sertoli cells of type B tubules (Frøland and Skakkebæk, 1971). Albeit the scRNAseq data also identified two types of Sertoli cells, with and without expression of *XIST* (Mahyari et al., 2021), the inherent lack of spatial information in scRNAseq experiments makes it impossible to link *XIST* expression in Sertoli cells to type A or B tubules.

We hypothesize that only Sertoli cells which lose the additional X-chromosome can mature properly and induce gametogenesis at puberty. To address this, we examined the expression of *XIST* during testicular development as well as the X-chromosome DNA ploidy in adult men with KS having both type A and B SCO tubules and focal spermatogenesis.

## Results

### Micro-mosaicism of type A and B Sertoli cells

To evaluate the expression of *XIST* in testes from men with KS, we performed RNA single-molecule *in situ* hybridization (smISH). To verify that *XIST* is expressed when two X-chromosomes are present, we stained an ovarian tumor which as expected showed a single red dot per nucleus (Fig. S1A) indicating expression of *XIST* and thereby X-inactivation. In testis biopsies from men with a normal testicular histology (Table S1), all somatic cell types were negative for *XIST*. However, spermatogonia and occasionally spermatocytes showed expression of *XIST* (Fig. S1B), which is in agreement with previous studies (Richler et al., 1992; Salido et al., 1992). In contrast, testis biopsies from patients with KS showed prominent expression of *XIST* in all somatic cell types, apart from some Sertoli cells (Fig. S1C and Fig. 1A). Furthermore, a testis biopsy from a man with a 46,XY karyotype but with a cellularity resembling KS (SCO tubules and Leydig cell hyperplasia) was negative for *XIST* expression (Fig. S1D). In total, testis biopsies from thirteen adult men with KS, all of which contained type A and B SCO tubules and with seven also containing areas with focal spermatogenesis, were investigated for *XIST* expression (Table S1). All KS biopsies contained SCO tubules with and without *XIST* expression (Fig. 1A, Fig. S2, and Table S1). Blinded histological evaluation by an experienced pathologist of serial sections stained with Hematoxylin and Eosin (n=6) confirmed that type A tubules were negative for *XIST*, whereas type B tubules were positive for *XIST*. Out of 270 tubules analyzed, 184 and 86 were classified as type A and B, respectively, with only six tubules classified as type A and expressing *XIST* (Fig. S3). After careful re-evaluation of the misclassified tubules, we found that the misclassified tubules contained a mixture of *XIST-*positive and -negative Sertoli cells, and that the *XIST* staining appeared different from the remaining type B Sertoli cells, with several dots per nucleus (Fig. S4), potentially indicating a transition phase where Sertoli cells are about to lose their *XIST* expression.

**Fig. 1:**
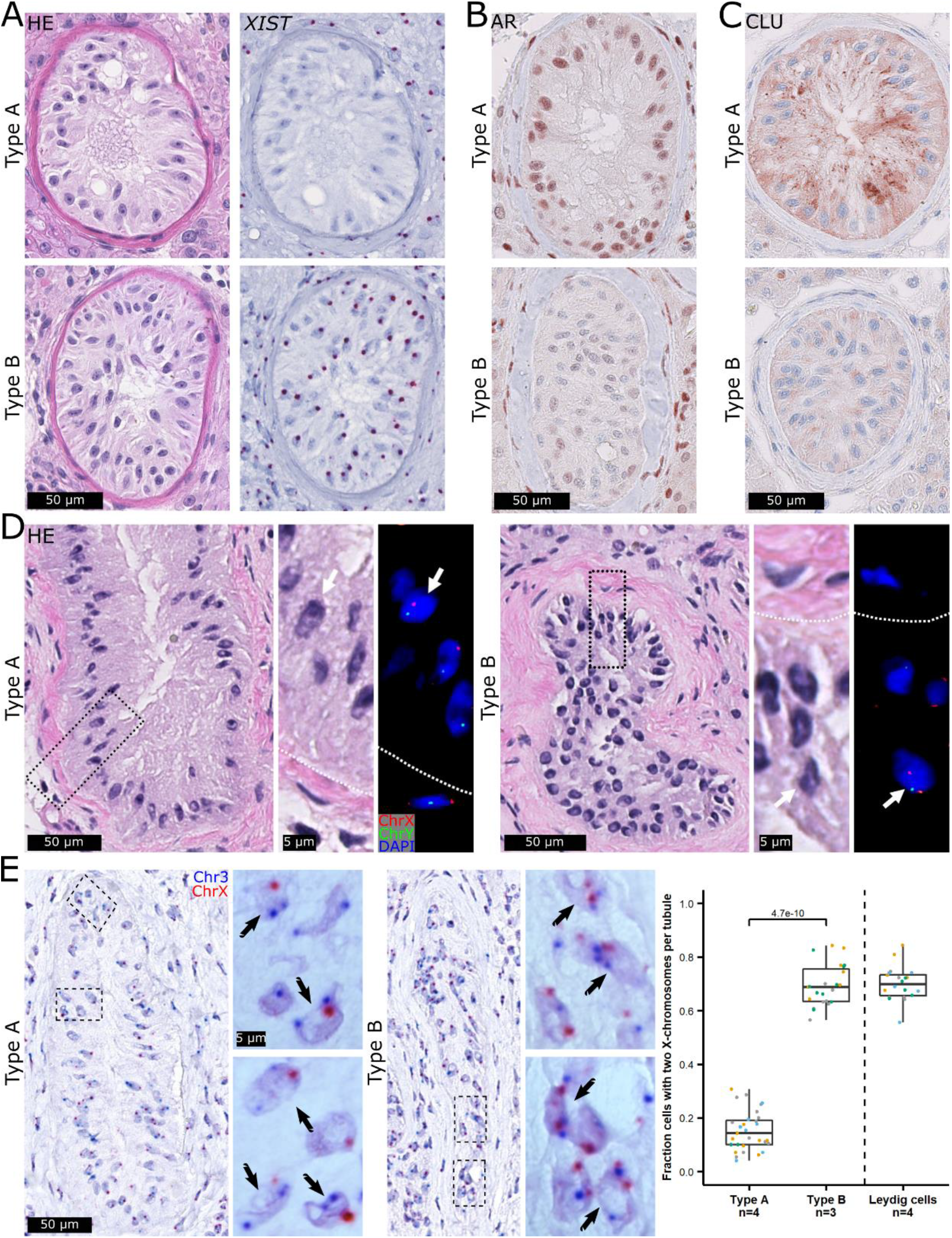
*XIST* expression differentiates mature type A and immature-like type B tubules and is correlated with the number of X-chromosomes. Sertoli cell-only (SCO) tubules classified as type A and type B on an HE staining were negative and positive for *XIST* RNA expression, respectively (**A**). Type A SCO tubules showed high protein levels of the Sertoli cell maturity markers androgen receptor (AR; **B**) and Clusterin (CLU; **C**), which was in contrast to type B tubules. DNA fluorescence *in situ* hybridization (FISH) against the X- (red) and Y-chromosome (green) (n=1) revealed a tendency towards type A tubules being 46,XY and type B being 47,XXY. (**D**). DNA single-molecular (sm)ISH against chromosomes 3 (blue) and X (red) (n=4) showed that type A tubules generally only contained one X-chromosome signal whereas type B tubules contained two (the tubules were identified on the basis of their nuclear morphology and *XIST* expression, see Fig. S6) (**E, left**). The mean fractions of Sertoli cells with two X-chromosome signals were around 15% in type A tubules (N= 31 tubules with a total of 887 Sertoli cells) and 69% in type B tubules (N=23 tubules with a total of 927 Sertoli cells) and statistically significantly different between the two types of tubules (p=4.7e-10, Wilcoxon Rank Sum Test). The fraction in type B tubules was similar to the one in Leydig cells (N=20 areas with a total of 1291 Leydig cells) serving as an internal control (**E, right**). Scalebars represent 50 and 5 µm on the low and high magnifications, respectively.

To investigate if Sertoli cells in type A tubules are mature as their nuclei morphology and localization along the basement membrane in the tubules suggest, we stained with antibodies against three Sertoli cell maturity markers, the androgen receptor (AR) (Sharpe et al., 2003; Wang et al., 2022), Clusterin (CLU, also known as SGP-2), which is secreted from mature Sertoli cells (Sharpe et al., 2003; Sylvester et al., 1984), and defensin beta 119 (DEFB119), a newly identified marker (Mahyari et al., 2023; Zhao et al., 2020). All three markers were highly expressed in type A Sertoli cells which was in contrast to type B Sertoli cells showing low expression levels (Fig. 1B,C and Fig. S5A) and to fetal Sertoli cells which were negative (Fig. S5B-D). The expression patterns of AR, CLU and DEFB119 hence support that Sertoli cells in type A tubules are mature, whereas type B Sertoli cells are not.

To determine if the absence of *XIST* expression in type A tubules was due to loss of X-inactivation or of the additional X-chromosome, we performed DNA fluorescence ISH (FISH) against the X- and Y-chromosome. This indicated that the additional X-chromosome was lost in type A tubules (Fig. 1D). It was, however, not possible to quantify the FISH signals adequately due to a lack of sensitivity. We therefore implemented an ultra-sensitive DNA smISH technique against chromosomes 3 and X (Fig. 1E, left). When applied to four biopsies, only a small fraction of the type A Sertoli cells had two X-chromosome signals (Fig. 1E, right) which was significantly lower than the high fractions in type B Sertoli cells and Leydig cells, serving as an internal control. Collectively, this suggests that Sertoli cells in type B tubules are not mature and have a 47,XXY karyotype, whereas the Sertoli cells in type A tubules are mature and have a 46,XY karyotype, and, therefore, that *XIST* expression is lost in type A mature Sertoli cells due to a micro-mosaic loss of the additional X-chromosome.

### Micro-mosaicism of spermatogonia and Sertoli cells supporting focal spermatogenesis

As in the 46,XY control (Fig. S1B), spermatogonia in focal spermatogenesis in KS were generally positive for *XIST* (Fig. 2A), whereby *XIST* expression cannot be used as a proxy for X-chromosome ploidy. But interestingly, as in type A tubules, Sertoli cells supporting focal spermatogenesis were generally devoid of *XIST* expression (Fig. 2A, Fig. S7A). DNA XY FISH on one biopsy showed a clear tendency towards Sertoli cells and spermatogonia having only a single X-chromosome (Fig. 2B). To confirm this, we used DNA smISH on two biopsies and observed that only small fractions of Sertoli cells and spermatogonia had two X-chromosome signals (Fig. 2C). These fractions were similar to the ones in type A tubules (Fig. 2D) and three controls (Fig. S7B and C), and very different from the ones in type B tubules and Leydig cells (Fig. 2D). Taken together, these data suggest that spermatogonia and Sertoli cells supporting focal spermatogenesis have a 46,XY karyotype.

**Fig. 2:**
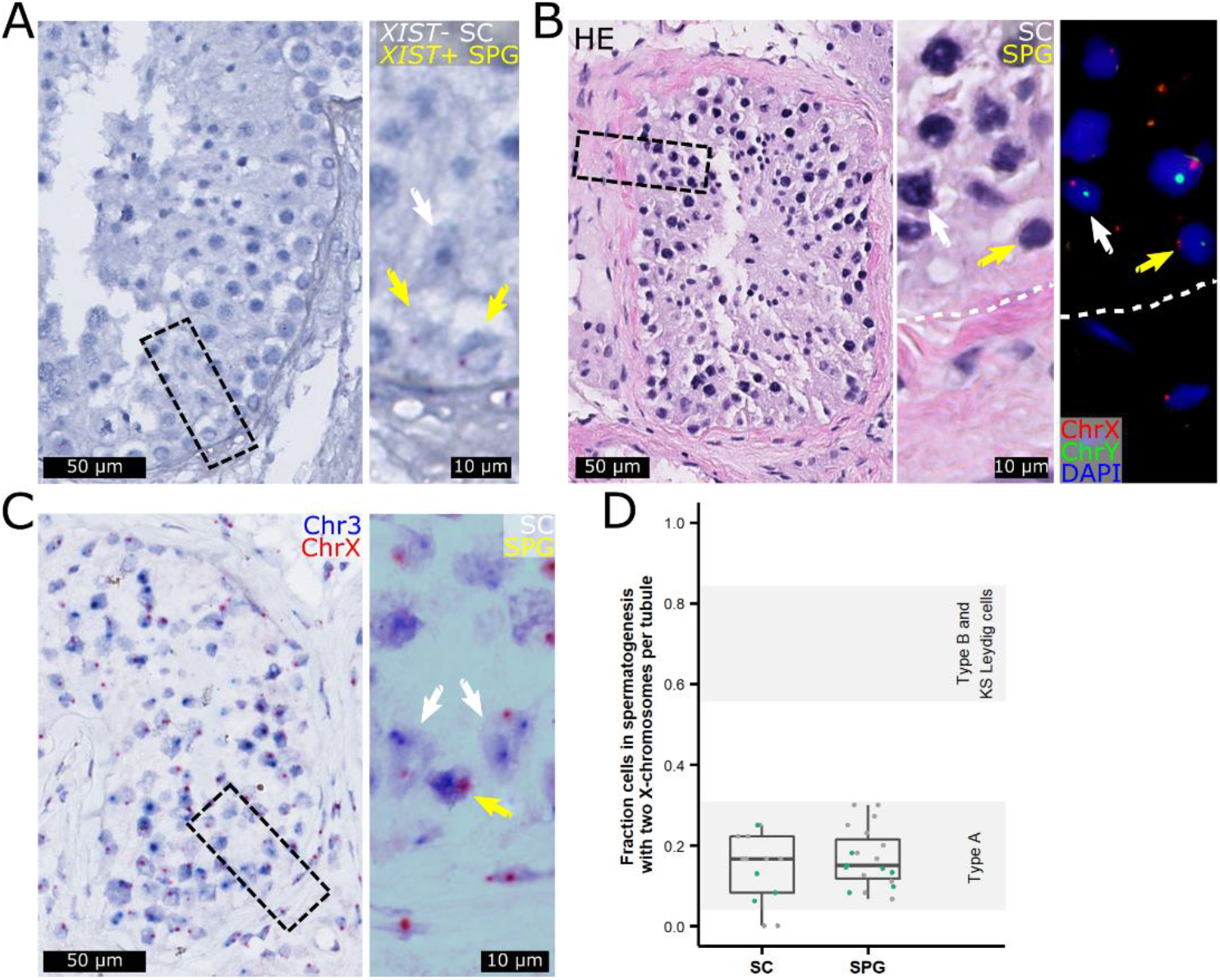
*XIST* is expressed in spermatogonia but is absent in Sertoli cells supporting focal spermatogenesis, and both cells are euploid. *XIST*-positive spermatogonia (SPG, yellow arrows) were present in testis specimens from men with Klinefelter syndrome, whereas the majority of the Sertoli cells (SC, white arrows) were *XIST*-negative (**A**). DNA FISH against the X- (red) and Y-chromosome (green) (n=1) revealed a tendency towards spermatogonia and Sertoli cells supporting spermatogenesis being 46,XY (**B**). DNA smISH against chromosomes 3 (blue) and X (red) (n=2) showed that spermatogonia and Sertoli cells generally only contained one X-chromosome signal (**C**). Scalebars represent 50 and 10 µm on the low and high magnifications, respectively. The mean fraction of spermatogonia (N=19 tubules with 299 spermatogonia) and Sertoli cells (N=13 tubules with 122 Sertoli cells) with two X-chromosome signals was around 15% and similar to type A tubules (lower shaded area) and very different from type B tubules and KS Leydig cells (upper shaded area) (**D**).

### Timing of X-chromosomal loss

Because loss of sex chromosomes in other settings shows an association to age (the number of cell divisions) (Leonova and Hanson, 1999; Pierre and Hoagland, 1972; United Kingdom Cancer Cytogenetics Group, 1992), we questioned whether loss of the additional X-chromosome in Sertoli cells of men with KS could be age-related and begin early in development. We therefore analyzed testis tissue from four fetuses with KS and one control aged gestational week (gw) 13 to 22. In all samples, the Leydig and Sertoli cells were generally positive for *XIST*, indicating that loss of *XIST* expression is not initiated in fetal life (Fig. 3A and Fig. S8). In the control, somatic cells were generally negative for *XIST* (Fig. S8). In all specimens, we observed a patchy expression of *XIST* in germ cells (Fig. 3A and S8).

**Fig. 3:**
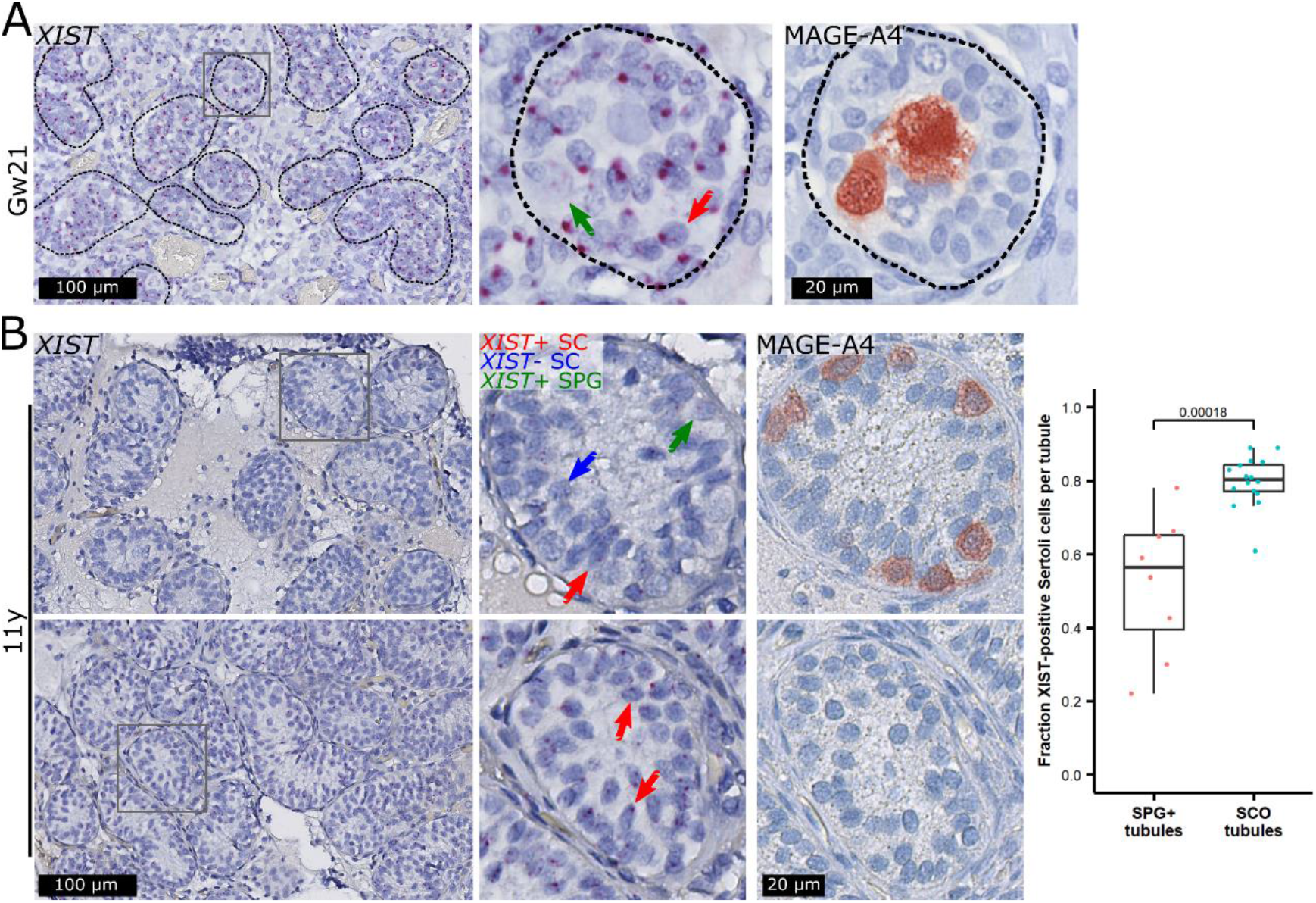
High *XIST* expression in Sertoli cells in fetuses but low *XIST* expression in tubules with spermatogonia compared to Sertoli cell-only (SCO) tubules in an 11-year-old prepubertal boy. In a gestational week (Gw) 21 fetus, *XIST* was expressed in virtually all Sertoli cells (SC, red arrow) but showed a patchy expression pattern in spermatogonia (green arrow, identified with a MAGE-A4 antibody) (**A**). The same pattern was seen in three additional fetuses (Fig. S8). In an 11-year-old prepubertal boy, spermatogonia (SPG) also showed a patchy expression pattern of *XIST* (**B, top**; green arrow). In tubules with spermatogonia only a fraction of Sertoli cells expressed *XIST* (red arrow) while the remaining were negative (blue arrow). This was in contrast to SCO tubules (**B, bottom**). Scalebars represent 100 and 20 µm on the low and high magnifications, respectively. The mean fractions of *XIST*-positive Sertoli cells were around 52% in SPG+ tubules (N=8 tubules with a total of 409 Sertoli cells) and 80% in SCO tubules (N=16 tubules with 1373 Sertoli cells) and statistically significantly different between the two types of tubules (p=0.00018, Wilcoxon Rank Sum Test) (**B, right**).

We further investigated testis biopsies from four boys with KS, which as expected (Aksglaede et al., 2006; Van Saen et al., 2018; Winge et al., 2020) contained few spermatogonia. In a biopsy with SCO from a nine-year-old prepubertal boy, the Sertoli cells were generally positive for *XIST* (Fig. S9, top). In a biopsy from an 11-year-old prepubertal boy, eight of the tubules contained spermatogonia, and in these, the fraction of Sertoli cells expressing *XIST* was significantly lower compared to tubules with SCO (Fig. 3B). In biopsies with SCO from two peri-pubertal boys aged 13 and 14 years, a small proportion of the tubules were *XIST*-negative and showed a morphology resembling type A Sertoli cells, while the remaining *XIST*-positive Sertoli cells resembled type B (Fig. S9, middle and bottom). Interestingly, in the biopsy from the 14-year-old boy, a high proportion of the tubules were either completely hyalinized or degenerating with few *XIST*-positive Sertoli cells (Fig. S9, bottom).

Taken together, these data show that fetal Sertoli cells as well as prepubertal Sertoli cells in SCO tubules express *XIST* and hence presumably have the 47,XXY karyotype. At puberty, *XIST*-positive Sertoli cells either resembled immature Sertoli cells or were present in degenerating tubules. On the other hand, Sertoli cells supporting spermatogonia in the prepubertal KS testis appear to gradually lose *XIST* expression and hence presumably the additional X-chromosome, while as the Sertoli cells mature in the peripubertal boys, even without the presence of germ cells, the X-chromosome will be lost as well.

## Discussion

The mechanism behind the partial testicular degeneration, and particularly the germ cell loss in men with KS has been debated for decades. Since men with KS generally father children with normal karyotypes (Lanfranco et al., 2004; Maiburg et al., 2012), it can be speculated whether germ cells can complete meiosis with an additional X-chromosome, which several studies have investigated albeit with conflicting results (Bergère et al., 2002; Blanco et al., 2001; Foresta et al., 1999; Garcia-Quevedo et al., 2011; Gonsalves et al., 2005; Lue et al., 2010; Sciurano et al., 2009; Yamamoto et al., 2002).

Our study focused on Sertoli cells for two reasons. First, the impairment of the Sertoli cell function has been increasingly recognized as the primary mechanism responsible for the pathogenesis of the testicular phenotype in patients with KS (Mahyari et al., 2021; Winge et al., 2018, 2020; Zhao et al., 2020). Secondly, the function of germ cells and the meiotic onset are physiologically controlled by somatic cells (Franca et al., 2016).

Our finding that both *XIST*-positive and -negative Sertoli cells can be found in testis biopsies from men with KS is in line with recent scRNAseq data (Mahyari et al., 2021), which is supported by the studies identifying the two types of SCO tubules (Skakkebaek, 1969) with Sertoli cells in type A tubules not having a Barr body (Frøland and Skakkebæk, 1971). In the latter studies, type A and B tubules were identified from morphological inspection, where their presumed maturity was accessed on the basis of their nuclear morphology. In this study, we show that the *XIST*-negative type A SCO tubules express the maturity markers AR, CLU, and DEFB119, whereas the expression levels of these markers in *XIST*-positive type B tubules were very low confirming the original morphological analysis. Using DNA FISH and DNA smISH, with the latter method found to be much more sensitive, we show that type A SCO tubules only have one X-chromosome whereas type B have two. To our knowledge, this is the first study using the combination of *XIST* RNA ISH and DNA ISH to investigate this. The only other study investigating the ploidy of Sertoli cells in SCO tubules is without spatial information, but rather using scRNAseq, where the authors conclude that lack of *XIST* expression is rather due to lack of X-inactivation (Mahyari et al., 2021). We also show that in adult men with KS, germ cells were only found inside tubules containing Sertoli cells with a nuclear morphology resembling type A Sertoli cells, lacking *XIST* expression, and with only one X-chromosome. And despite expressing *XIST*, the spermatogonia also had a single X-chromosome. The focus of previous studies using DNA XY FISH was to access the ploidy of germ cells in tubules with spermatogenesis. Apart from one study where only ten spermatogonia were counted (Foresta et al., 1999), previous studies have not distinguished spermatogonia from primary spermatocytes in their ploidy analysis (Bergère et al., 2002; Blanco et al., 2001; Garcia-Quevedo et al., 2011; Gonsalves et al., 2005; Sciurano et al., 2009; Yamamoto et al., 2002), which makes it difficult to compare the results with ours. In four of the studies, the ploidy of Sertoli cells was also analyzed. Two studies found that all Sertoli cells were 47,XXY (Foresta et al., 1999; Yamamoto et al., 2002), one study found a high fraction of Sertoli cells to be 47,XXY (Sciurano et al., 2009), whereas the last study found that around half of the Sertoli cells were 47,XXY (Garcia-Quevedo et al., 2011). Three of the studies used spread cells meaning that the testicular architecture was not preserved (Foresta et al., 1999; Garcia-Quevedo et al., 2011; Sciurano et al., 2009), whereas the last study was performed on tissue fragments implying that the testicular architecture was only somewhat intact (Yamamoto et al., 2002). It is hence difficult to know whether the investigated Sertoli cells represent Sertoli cells of type A or B SCO tubules.

Loss of the sex chromosomes has been documented mainly for the Y-chromosome (Jacobs et al., 1963; Pierre and Hoagland, 1972) and occurs presumably during cell division. The timing of the X-chromosome loss is difficult to establish because mitotic cell divisions occur in Sertoli cells and germ cells throughout testis development. In addition, 46,XY germ cells express *XIST* and retain transcriptional activity of numerous X-chromosomal genes (Wang et al., 2001) suggestive of an X-inactivation that is different from the one in somatic cells. Nevertheless, our study provides some novel observations concerning the timing of the presumed ‘correction’ of aneuploidy in Sertoli cells, which appears to occur postnatally, most likely at the onset of puberty, concurrently with the activation of Sertoli cell proliferation. Whether the loss of the additional X-chromosome in the spermatogonia also happens at puberty, and in which cell type the additional X-chromosome is lost first cannot be determined from our data as we were not able to perform DNA smISH on fetal and prepubertal KS samples. Hence, we cannot rule out that micro-mosaicism of germ cells could originate already during early embryonic development.

Based on our data, we propose a model which stipulates that the remarkable heterogeneous testicular phenotype seen in adult men with KS is due to micro-mosaicism for the X-chromosome aneuploidy in the Sertoli cells and germ cells (Fig. 4). Until puberty, the histological pattern is rather uniform - all tubules are preserved, although the number of spermatogonia is severely reduced (Aksglaede et al., 2006; Van Saen et al., 2018; Winge et al., 2020). During puberty, when differentiation of Sertoli cells and adult spermatogenesis are induced by activation of the hypothalamic–pituitary–gonadal (HPG) axis, the fate of the Sertoli cells will have four scenarios dependent on their ploidy and the presence of germ cells in the tubules. In the tubules devoid of germ cells, it is only euploid Sertoli cells that are able to mature hence forming type A SCO tubules, whereas the aneuploid Sertoli cells are unable to mature thus causing them to arrest in the immature-like type B state. If the aneuploid Sertoli cells attempt to differentiate, they will fail causing extensive cell death and degeneration, with subsequent hyalinization and the appearance of “ghost” tubules. In rare tubules with surviving spermatogonia, mature euploid Sertoli cells can induce maturation of germ cells, and only the subset of spermatogonia that eliminated the supernumerary X-chromosome would become able to form foci of complete spermatogenesis.

**Fig. 4:**
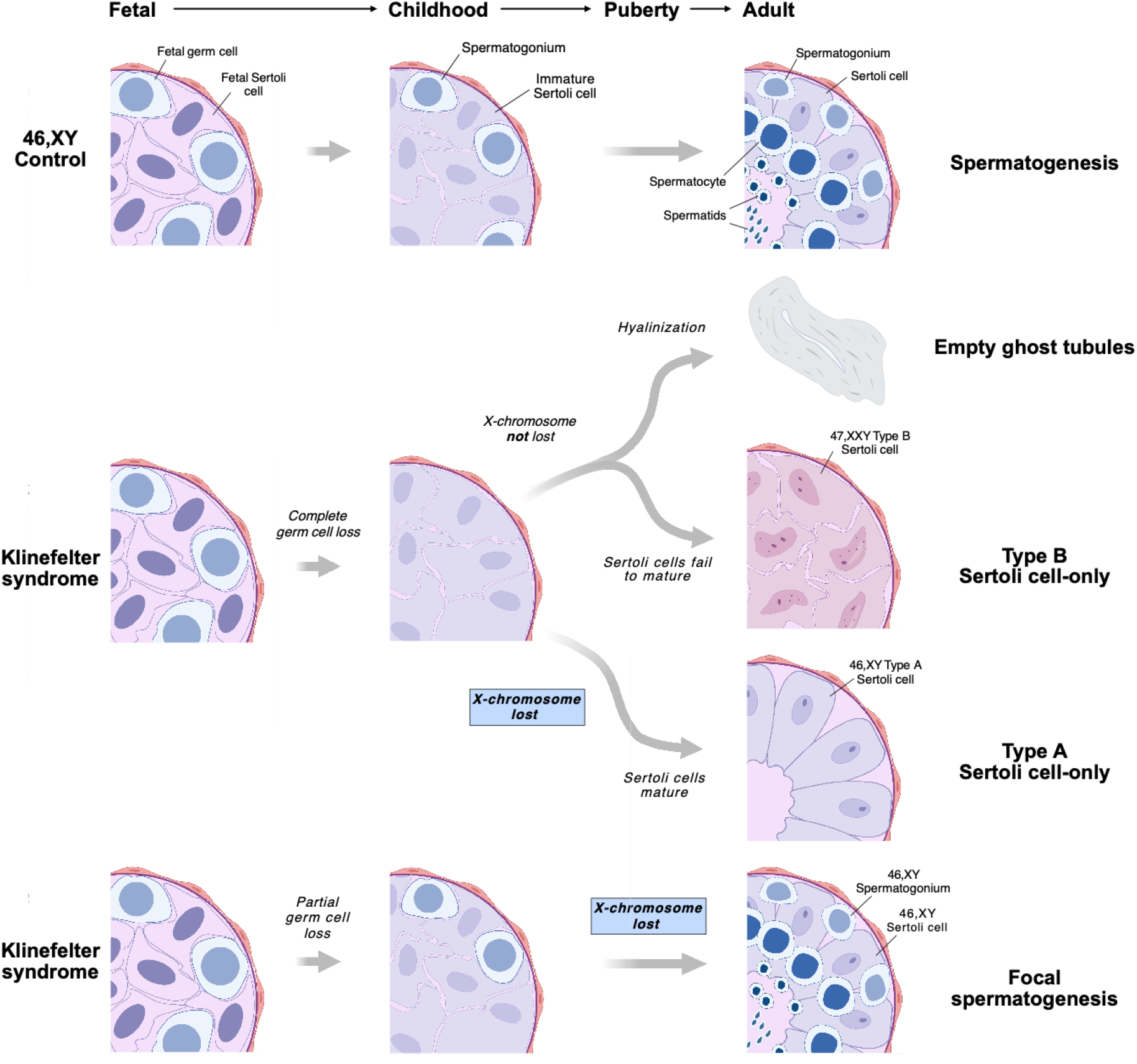
Schematic illustration of germ cell loss and the proposed model of X-chromosome loss in specific cell stages in the testes from men with Klinefelter syndrome (KS). During fetal life (first column), the seminiferous tubules contain fetal germ cells (initially gonocytes and later pre-spermatogonia) and fetal Sertoli cells both in the 46,XY control (first row) and in KS (second and third row). In controls, the tubules contain spermatogonia and immature Sertoli cells in childhood (second column). Puberty leads to initiation of spermatogenesis (third column). For KS, testicular pathology is changing during childhood, puberty and adulthood leading to either a complete or partial loss of germ cells. Depending on the presence of germ cells and the ploidy of the Sertoli cells, four possible scenarios exist. The first three scenarios involve a complete loss of germ cells during childhood. In the first scenario, the Sertoli cells will attempt to mature leading to cell death and hyalinized tubules. In the second scenario, the Sertoli cells fail to mature thus creating morphologically distinct type B tubules. In the third scenario, a loss of the additional X-chromosome will lead to Sertoli cell maturation forming type A tubules. The fourth scenario implies that only a partial germ cell loss has occurred during childhood, and if the Sertoli cells and the spermatogonia lose the additional X-chromosome, focal spermatogenesis will occur. The figure is for illustration purposes and the actual order of events as well as the number of germ cells may be different. Figure created with BioRender.com.

This study builds on different types of histological staining. A challenge when using such staining is that either the signal can be out of the sectioning plane, resulting in false negative cells, or the signal in a given nucleus originates from another cell on top of the quantified cell resulting in a false positive cell. To confirm that our findings were true and not merely a result of high numbers of false positive or negative cells, we compared our quantifications in KS to 46,XY controls, and found no differences. Spatial transcriptomics/proteomics or stereological analyses would be needed to validate our findings, but in general we believe the technical bias in our study is low.

Our findings have important clinical implications, especially in view of the increasing use of assisted reproductive techniques helping men with KS to fulfil their wish of biological paternity. First, our results reassure that the risk of fathering a child with abnormal X-chromosome ploidy is similar to men with a 46,XY karyotype, which is in agreement with other studies (reviewed in Maiburg et al., 2012). Second, since it appears that only euploid spermatogonia are present in adulthood, it suggests that the loss of the additional X-chromosome in spermatogonia does not occur gradually after puberty, which is in line with a comprehensive meta-analysis showing that the TESE success rate is not affected by age (Corona et al., 2017). Lastly, due to the requirement of micro-mosaicism for normal sex chromosome ploidy simultaneously in Sertoli cells and the adjacent spermatogonia which according to our data happens at puberty, it may be unwarranted to cryopreserve testicular tissue from children with KS for later transplantation, which is in line with previous research (reviewed in Shepherd and Oates, 2021).

## Materials and methods

### Samples

The use of tissue was approved by a regional medical and research ethics committee (H-2–2014-103) and the Danish Data Protection Agency (2012–58-0004, local no. 30–1482, I-Suite 03696) as well as the ethics committee of the University Hospital in Brussels (UZ Brussel), Belgium (2015/121; BUN 143201524312). Sample descriptions can be found in Table S1. Testis tissue from four fetuses were collected between 1993 and 2012 at autopsy of induced abortions due to KS of the fetus (only samples with 47,XXY or 48,XXYY karyotypes were included). Two additional samples from control fetuses were included as well. Gestational age varied between 13 and 22 weeks. Samples from adult patients were collected between 1973 and 2022. 14 testis samples from adult men with KS aged 18 to 32 years were included. In addition, tissue from eight adult controls were included. These comprised five samples that were obtained from orchiectomy specimens from individuals with testicular cancer, a specimen obtained for testicular sperm extraction (TESE) due to obstructive azoospermia, a specimen taken due to infertility of the patient, as well as a specimen from an ovarian tumor. Four testis samples from boys with KS aged 9 to 14 years were collected between 1976 and 1983. Following surgical excision, the samples were immediately fixed in different fixatives: 10% neutral buffered formalin (NBF), 4% paraformaldehyde (PFA) in PBS, acidified formal alcohol fixative (AFA), Bouin’s fixative, GR-fixative (7.4% formaldehyde, 4% acetic acid, 2% methanol, 0.57% sodium phosphate, dibasic and 0.11% potassium phosphate, monobasic), Stieve’s fixative, or Cleland’s fixative for at least 16 hours. The samples were then dehydrated and embedded in paraffin.

### RNA single-molecule in situ hybridization and quantification

The RNA single-molecule *in situ* hybridization (smISH) experiments were performed on 4 μm sections mounted on SuperFrost Plus™ Slides (ThermoFisher Scientific, MS, USA) using the RNAscope® 2.5 HD Detection Reagent - RED kit according to the manufacturer’s recommendations (Advanced Cell Diagnostics, CA, USA). Briefly, testicular tissue sections were dewaxed in xylene and washed in 100% ethanol followed by treatment with hydrogen peroxide for 10 min. Target retrieval was performed for 15 min at 99 °C followed by treatment with protease plus for 30 min at 40 °C. The slides were hybridized with the *XIST* probe (Cat No. 311231) for 2 hours at 40 °C followed by a series of signal amplifications (with amplification round 5 for 30 min (NBF, AFA, PFA, Bouin’s and GR-fixative) or 60 min (Stieve’s and Cleland)). The sections were counterstained with Mayer’s hematoxylin and dried before mounting with Vectamount® Permanent Mounting Medium (Vector Laboratories, CA, USA). The negative control probe *DapB* (a bacterial RNA; Cat No. 310043) was run in parallel with the *XIST* probe and showed less than 10% positive cells.

Sertoli cells in tubules with spermatogenesis were identified based on nuclei morphology. Only tubules where at least six Sertoli cells could be identified were counted. A total of 41 tubules from three controls, aNorm1-3, (a total of 766 Sertoli cells) and all tubules from KS patients (N=34 tubules with 467 Sertoli cells) were counted as being positive or negative for *XIST* with no discrimination in dot intensity being made. Sertoli cells in a testicular biopsy from a prepubertal boy with KS (pKS2) were identified based on nuclei morphology and MAGE-A4-negativity. A total of eight tubules also containing spermatogonia (409 Sertoli cells) and 16 Sertoli cell-only (SCO) tubules (1373 Sertoli cells) were counted as being positive or negative for *XIST* with no discrimination in dot intensity being made.

### DNA in situ hybridization and quantification

DNA ISH staining was only compatible with formalin-fixed tissue, and hence only performed on specimens fixed in NBF, PFA, and AFA (Table S1).

DNA fluorescence ISH (FISH) was performed on a 2 μm section from aKS7 (Table S1) using the Zyto*Light* FISH-Tissue Implementation Kit according to the manufacturer’s recommendations (Zytovision GmbH, Germany). Briefly, the slide was heated to 70 °C for 10 min, followed by dewaxing in xylene and rehydrated through an ethanol series to water. Target retrieval was performed at 98 °C for 15 min and the slide was incubated in a Pepsin Solution for 15 min at 37 °C. After heating the slide to 70 °C for 10 min, the slide was incubated overnight at 37 °C with probes targeting the α satellites of the X-chromosome (DXZ1; Xp11.1-q11.1, red color) and satellite 3 on the Y-chromosome (DYZ1; Yq12, green color). The slide was dehydrated by an ethanol series and dried before mounting with DAPI/DuraTect-Solution. The section was visualized at a 100X magnification using an Olympus BX61 microscope and captured using the Cell Sense Dimensions V1.6 software (Olympus Ltd., Denmark). Type A and type B SCO tubule classification was performed on a 2 µm serial section stained with Hematoxylin and Eosin. Due to the low number of cells positive for both probes, no quantifications were made.

DNA smISH with chromogen detection was performed on 4 µm sections using the DNAscope™ HD Duplex Detection Kit according to the manufacturer’s recommendations (Advanced Cell Diagnostics). Briefly, the slides were dewaxed in xylene and washed in 100% ethanol followed by treatment with hydrogen peroxide for 10 min at room temperature and protease plus for 20 min at 40 °C. Target retrieval was performed for 30 min at 99 °C. The slides were hybridized with the DS-Hs-CEPX-q-C1 probe (Cat. No. 1121921-C1; Xq11.2; red color) and DS-Hs-CEP3q-C2 probe (Cat. No. 1080211-C2; 3q11.1-q11.2; blue color) overnight at 40 °C followed by a series of signal amplifications. The sections were counterstained with Mayer’s hematoxylin (NBF- and PFA-fixed specimens) or Gill’s hematoxylin (AFA-fixed specimens) and dried before mounting with Vectamount® Permanent Mounting Medium.

Quantifications of X-chromosome content was only performed on cells having at least one blue and one red dot, and the fraction of cells having two X-chromosome signals was only calculated for tubules having at least six quantifiable cells. Type A (N=31 tubules from four biopsies) and B (N=23 tubules from three biopsies) SCO tubules as well as Leydig cells (N=5 areas from each of four biopsies) were identified based on morphology. A total of 887, 927, and 1291 type A Sertoli, type B Sertoli, and Leydig cells were counted, respectively. Two of the four biopsies contained tubules with spermatogenesis, and in these, Sertoli cells were identified based on nuclei morphology and position in the tubules. A total of 122 Sertoli cells in nine tubules from aKS7 and four tubules from aKS8 as well as a total of 295 Sertoli cells in eight tubules from each of aNorm4 and aNorm5, as well as 14 tubules from aNorm6 were counted. Spermatogonia were identified based on morphology and position in the tubules. A total of 299 spermatogonia in 12 tubules from aKS7 and seven tubules from aKS8 as well as a total of 623 spermatogonia in 12 tubules from each of aNorm4, aNorm5, and aNorm6 were counted.

All slides were visualized using a NanoZoomer 2.0HT slide scanner (Hamamatsu Photonics, Japan).

The data was plotted in R v.3.6.1 using the packages ggplot2 (Wickham, 2016) and ggpubr (Kassambara, 2020). A Wilcoxon Rank Sum Test was used to test for statistically significant difference between the groups.

### Classification of type A and type B SCO tubules

An experienced pathologist (NES) evaluated a tissue section stained with Hematoxylin and Eosin adjacent to the one stained with the *XIST* probe by smISH and categorized the SCO tubules as either type A or type B. This was performed on six specimens with a total of 270 tubules evaluated (Fig. S3). Another observer (SBW) evaluated the section stained with the *XIST* probe and categorized the tubules as either being positive or negative for *XIST* (with less than 10% *XIST*-positive cells). A pairwise comparison was then performed.

### Immunohistochemistry

The IHC staining was conducted according to a standard indirect peroxidase method, as previously described (Nielsen et al., 2019). Briefly, tissue sections were subjected to heat-induced antigen retrieval in a pressure cooker (medical decloaking chamber, Biocare, Concord, CA, USA) either in 0.01 M citrate buffer (pH 7.4) or in TEG buffer (10 mM Tris, 1 mM EDTA, 0.05% Tween 20, pH 8.5) at 110 °C for 30 minutes. Endogenous peroxidase was blocked with 1% (v/v) H_2_O_2_ in methanol for 30 min. Unspecific staining was blocked using 0.5% skimmed milk in Tris-buffered saline (TBS) for 30 min. Sections were incubated overnight at +4 °C in a humidified chamber with primary antibodies diluted in TBS. Primary antibodies were MAGE-A4 (kindly provided by G. C. Spagnoli, University of Basel, Switzerland, Clone 57B, produced in mouse) diluted 1:1000, AR (cat. no. sc-816, Santa Cruz Biotechnology, TX, US, produced in rabbit) diluted 1:100, CLU (cat. no. MAB2937, R&D Systems, MN, US, produced in mouse) diluted 1:250, or DEFB119 (cat. no. PA5-60222, ThermoFisher Scientific, produced in rabbit) diluted 1:40. On the next morning, the slides were left at room temperature for 1 hour and then incubated for 30 min with the anti-mouse or anti-rabbit ImmPRESS HRP (peroxidase) secondary antibodies (Vector laboratories, CA, USA). Between all steps (except after the blockage of unspecific staining) the sections were washed in TBS. Visualization was performed using ImmPACT AEC peroxidase (HRP) substrate (Vector Laboratories). The sections were subsequently counterstained with Mayer’s hematoxylin and mounted with Aquatex® mounting medium (Merck KGaA, Germany).

## Acknowledgments

We thank the Novo Nordisk Foundation (grant no. NNF21OC0069913 to KA) for financial support. The authors wish to acknowledge Lisa Leth Maroun and Svetlana Teplaia for assistance with the DNA FISH staining, and John E. Nielsen and Ana Ricci C. G. Nielsen for technical assistance with the IHC experiments. We also want to thank Professor Mikkel H. Schierup for helpful input to the manuscript.

## Author contributions

SBW, NES, and KA conceived and designed the study. EG and LA provided samples. SBW established the smRNA and smDNA ISH methods and performed all experiments. SBW, NES, and KA did the experimental analysis. GS designed Figure 4. NES, ER-DM, AJ, and KA supervised the study. SBW and KA wrote the manuscript with input from all authors. All authors provided useful feedback and discussions.

