## Supplementary figures and table for "X-chromosome loss rescues Sertoli cell maturation and spermatogenesis in Klinefelter syndrome"

### Supplementary figures and tables

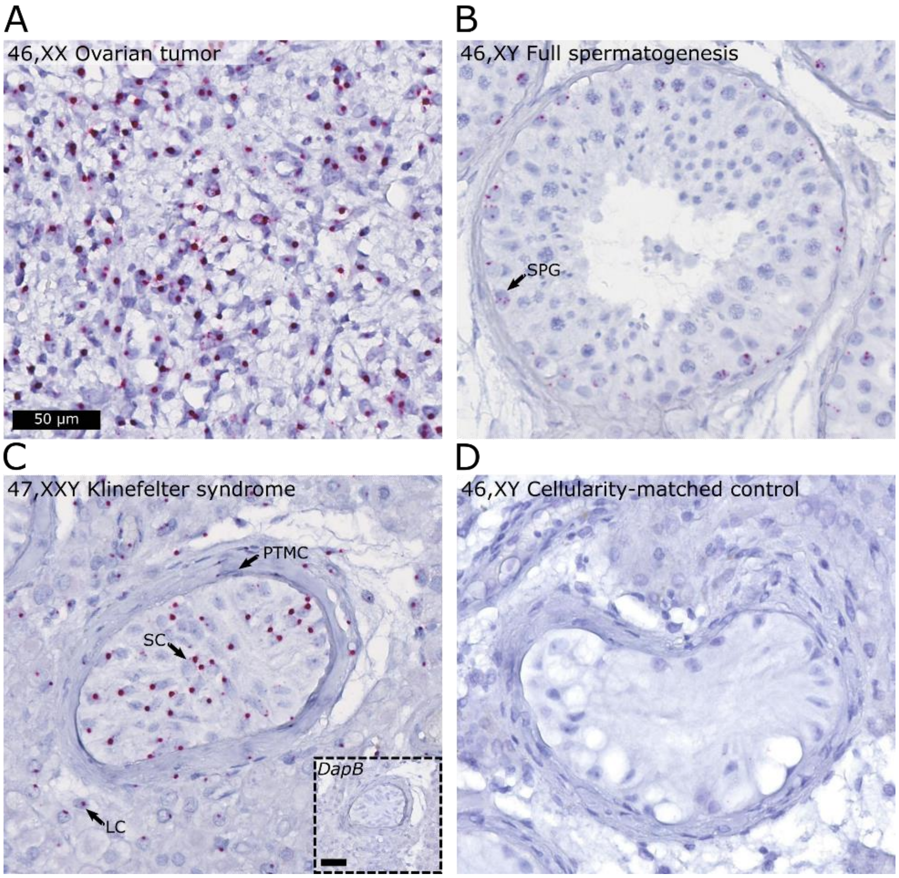


**Fig. S1: *XIST* is expressed in an ovarian tumor, spermatogonia of a normal control testis as well as in the somatic cells in a testis specimen from a man with Klinefelter syndrome (KS).** RNA single-molecule *in situ* hybridization (smISH) with a probe targeting the *XIST* transcript was performed on an ovarian tumor showing *XIST* expression in around 80% of cells (**A**), a testis biopsy from a normal man with spermatogenesis showing expression in spermatogonia (SPG) and occasionally spermatocytes (**B**), a testis biopsy from a man with KS showing expression in Leydig- (LC), peritubular myoid- (PTMC), and Sertoli cells (SC) (the inset shows staining with a probe targeting the bacterial RNA, *DapB*, which was negative) (**C**), and a testis biopsy from a 46,XY man with a cellularity-matched testicular histology (Sertoli cell-only pattern and Leydig cell hyperplasia) showing no *XIST* expression (**D**). Scalebars represent 50 µm.

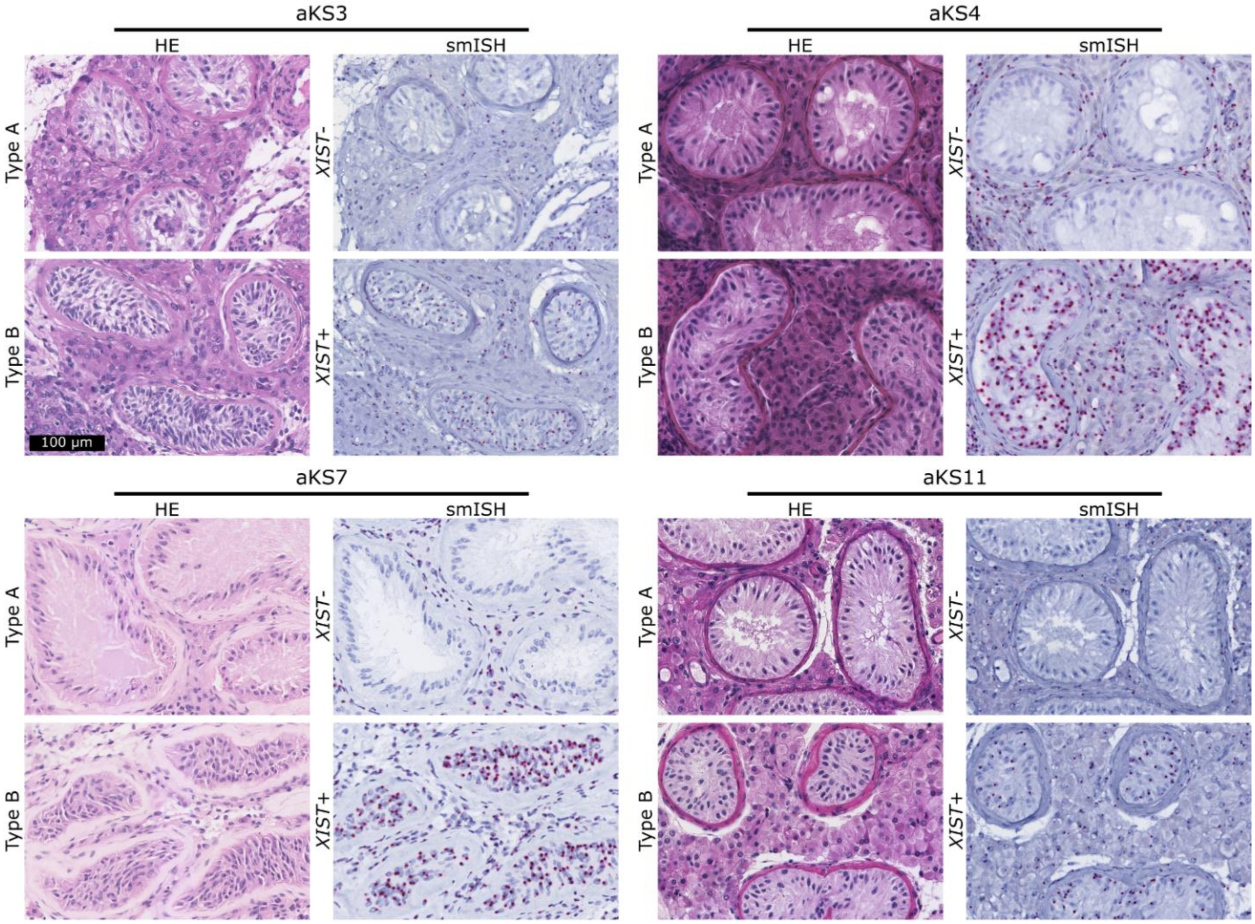


**Fig. S2. *XIST* expression differentiates type A and B Sertoli-cell only (SCO) tubules.** SCO tubules classified as type A and type B from Hematoxylin and Eosin (HE) staining of testicular biopsies from four men with Klinefelter syndrome (KS) (see Table S1 for descriptions of the samples). RNA single-molecule *in situ* hybridization (smISH) with a probe targeting *XIST* on a serial section showing a differential expression pattern of *XIST* with tubules classified as type A being negative for *XIST* and type B tubules being positive for *XIST*. A similar pattern was observed in a total of thirteen independent biopsies. Scalebar represents 100 µm.

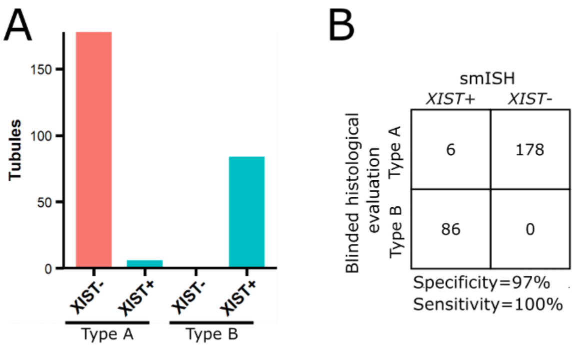


C

| Code | Type A and *XIST*-negative | Type A and *XIST*-positive | Type B and *XIST*-positive | Type B and *XIST*-negative |
|---|---|---|---|---|
| aKS3 | 14 | 3 | 12 | 0 |
| aKS4 | 10 | 0 | 9 | 0 |
| aKS7 | 28 | 0 | 7 | 0 |
| aKS9 | 8 | 0 | 14 | 0 |
| aKS11 | 105 | 3 | 40 | 0 |
| aKS13 | 13 | 0 | 4 | 0 |

**Fig. S3. Correlation between type A and B Sertoli cell-only (SCO) classifications and *XIST* expression per sample.** An experienced pathologist (NES) evaluated a tissue section stained with Hematoxylin and Eosin adjacent to the one stained with the *XIST* probe and categorized the SCO tubules as either type A or type B. This was performed on six specimens with a total of 270 tubules evaluated. All tubules (N=86) classified as type B were positive for *XIST* whereas 178 out of 184 classified as type A were negative for *XIST* (**A**). This yields a specificity of 97% and a sensitivity of 100% (**B**). The six tubules classified as type A but being positive for *XIST* came from two different biopsies (**C**).

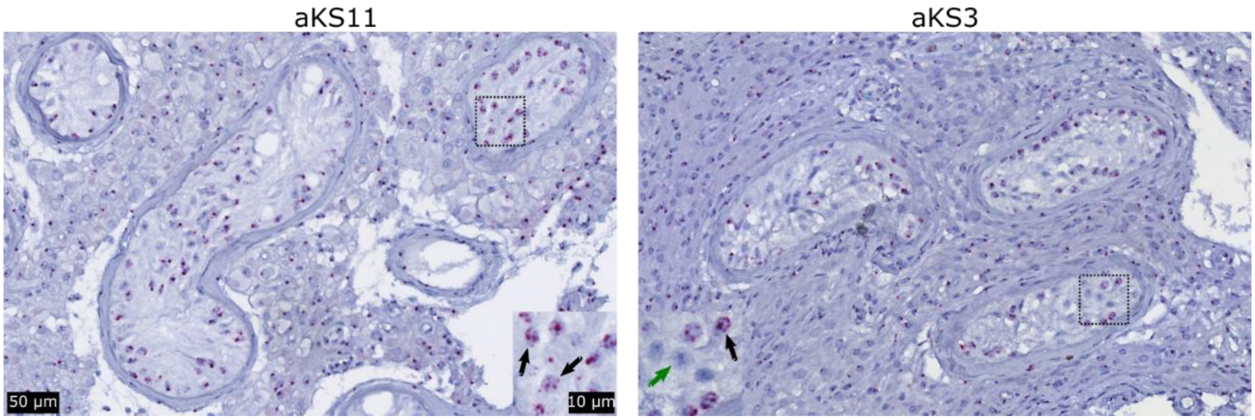


**Fig. S4: *XIST*-positive tubules classified as type A show distinct *XIST* expression patterns with more dots per cell and many negative cells.** A total of six Sertoli cell-only (SCO) tubules from two patients, aKS3 and aKS11 (see Fig. S3C) were classified on the basis of the Hematoxylin and Eosin (HE) staining as type A SCO tubules but were, in contrast to the remaining classified type A tubules (Fig. 1A and Fig. S2), positive for *XIST* expression. In these six tubules, the *XIST* staining generally showed more than one dot per nucleus (black arrows) in contrast to the general pattern with only one red dot per nucleus (see Fig. S1 and S2 for examples), and in aKS3, about half of the Sertoli cells were negative for *XIST* expression (green arrow). Scalebars represent 50 µm and 10 µm on the low and high magnifications, respectively.

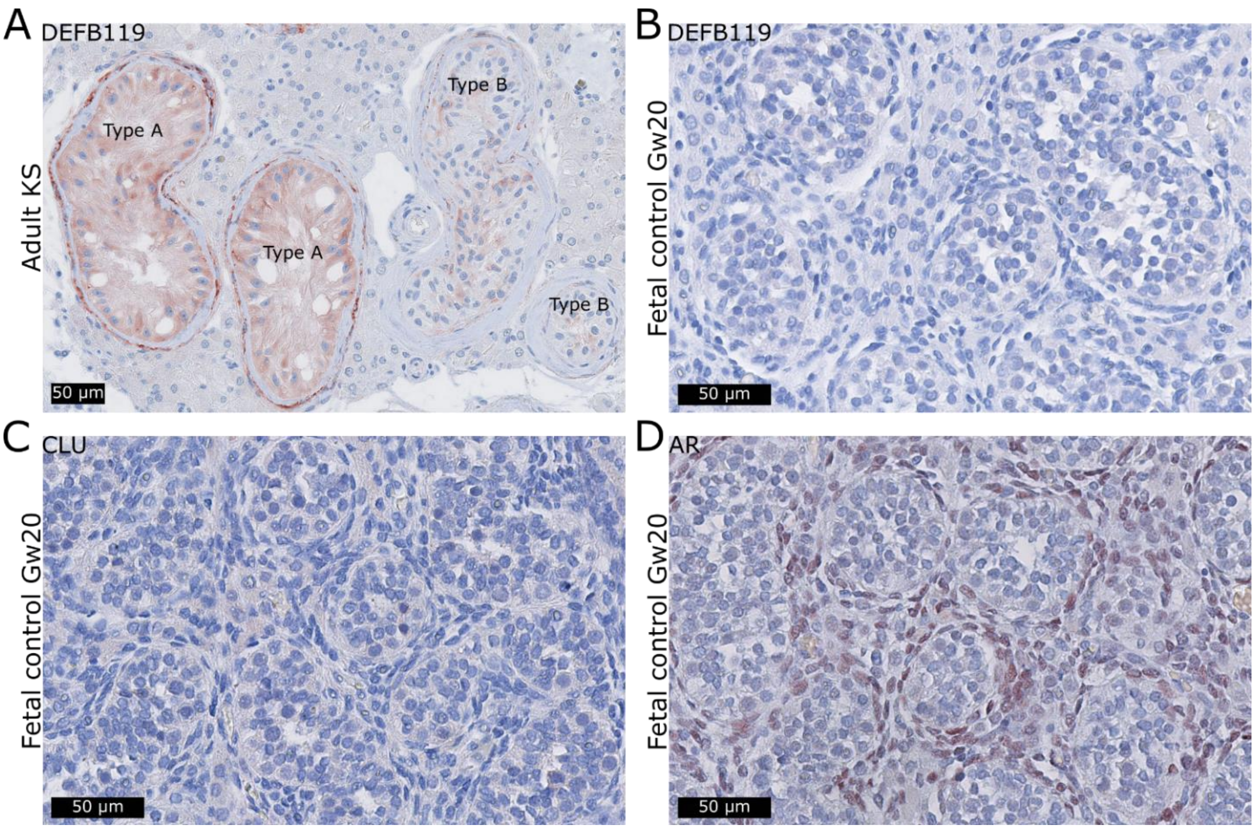


**Fig. S5: Expression of DEFB119 is high in type A and low in type B tubules, and AR, CLU and DEFB119 are not expressed in fetal Sertoli cells.** Immunohistochemical staining with an antibody against DEFB119 on a testis from an adult patient with Klinefelter syndrome (KS) showing that DEFB119 was highly expressed in type A tubules which was in contrast to type B tubules showing low expression level (**A**). In a testis specimen from a fetal control aged gestational week (Gw) 20, neither DEFB119 (**B**), CLU (**C**) or AR (**D**) were expressed in fetal Sertoli cells thus confirming their use as markers of mature Sertoli cells. AR was expressed by interstitial cells (**D**). Scalebars represent 50 µm.

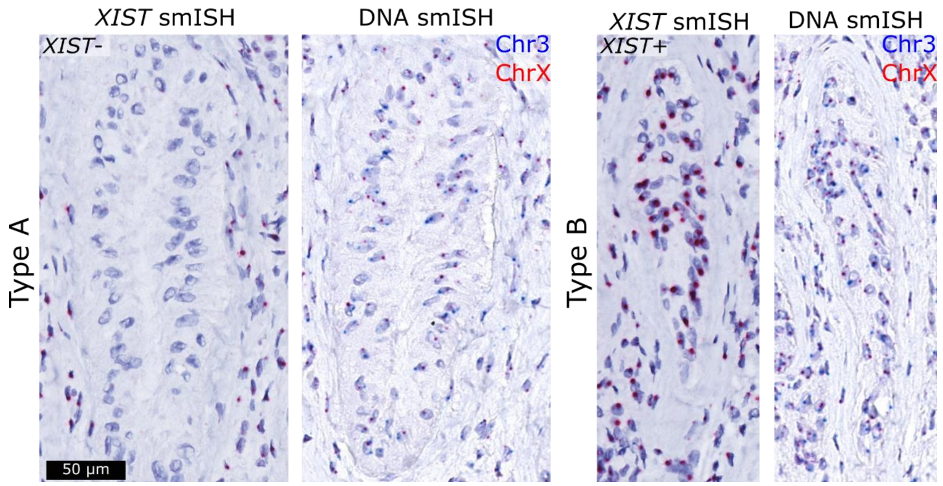


**Fig. S6: *XIST* differentiates type A and B Sertoli cell-only (SCO) tubules classified on the slides used for DNA single-molecule *in situ* hybridization (smISH).** Example of a type A SCO tubule classified morphologically from the DNA smISH slide showing no expression of *XIST* detected by RNA smISH, whereas type B SCO tubules showed *XIST* staining. The *XIST*-stained slide is four sections apart from the DNA-stained slide and hence, the tubules look a bit different. Quantification of the DNA smISH staining can be seen in Fig. 1E. Scalebar represents 50 μm.

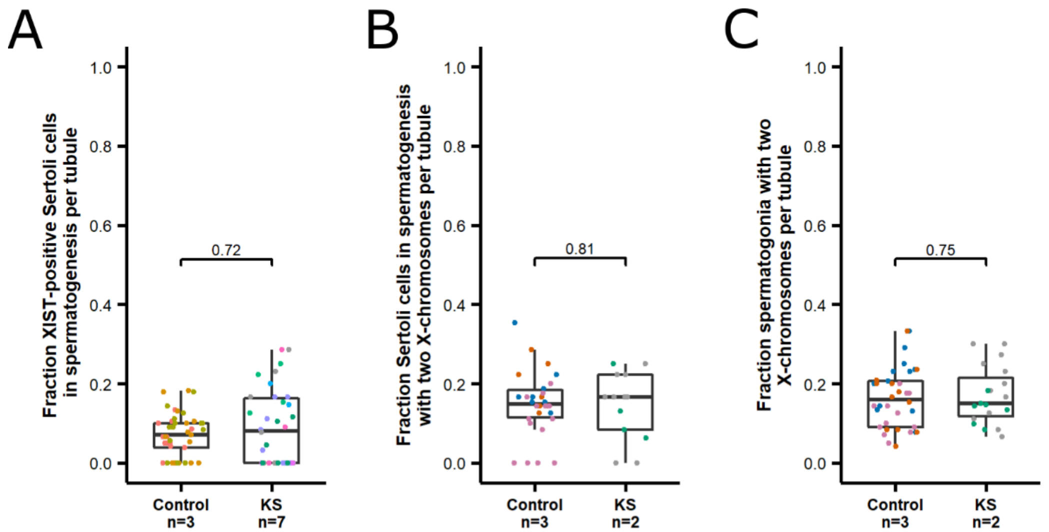


**Fig. S7: *XIST* expression and X-chromosome ploidy is not different between testicular biopsies from men with Klinefelter syndrome (KS) and controls.** RNA single-molecule *in situ* hybridization (smISH) against *XIST* showed that in tubules with spermatogenesis, the mean fraction of *XIST*-positive Sertoli cells was around 10% in both controls (n=3 samples with N=41 tubules and a total of 766 Sertoli cells) and KS (n=7 samples with N=34 tubules and a total of 467 Sertoli cells) and not statistically different between the two groups (p=0.72, Wilcoxon Rank Sum Test) (**A**). DNA chromogen smISH against chromosomes 3 and X showed that the mean fraction of Sertoli cells with two X-chromosome signals was around 15% in both controls (n=3 samples with N=30 tubules and a total of 295 Sertoli cells) and KS (n=2 samples with N=13 tubules and a total of 122 Sertoli cells) and not statistically significantly different between the two groups (p=0.81, Wilcoxon Rank Sum Test) (**B**). The mean fraction of spermatogonia with two X-chromosome signals was around 16% in both controls (n= 3 samples with N=36 tubules and a total of 623 spermatogonia) and KS (n=2 samples with N=19 tubules and a total of 299 spermatogonia) and not statistically significantly different between the two groups (p=0.75, Wilcoxon Rank Sum Test) (**C**).

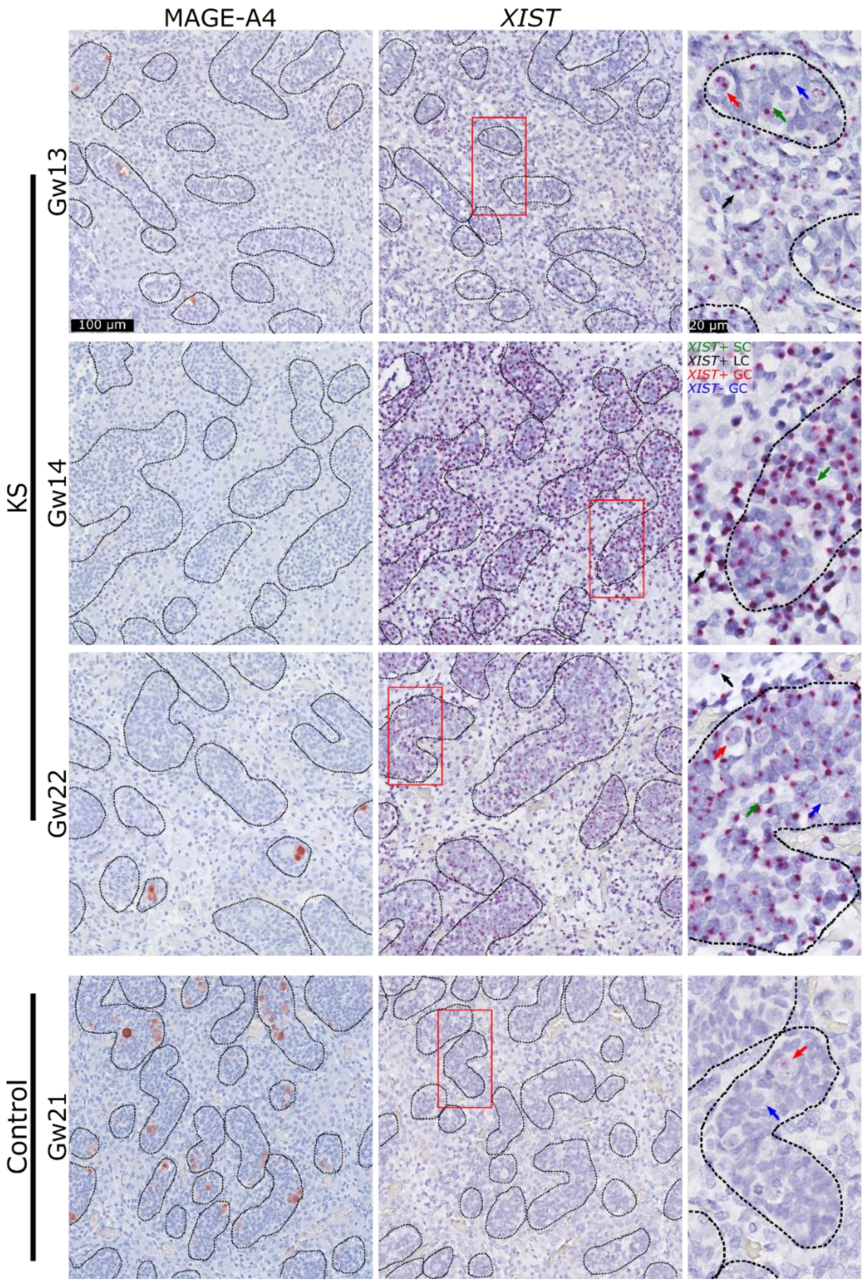


**Fig. S8: During second trimester, Leydig and Sertoli cells express *XIST* whereas the expression in germ cells is sporadic**. Three fetuses with Klinefelter syndrome (KS) and one control (aged gestational week (Gw) 13-22) were stained with the *XIST* probe (middle and right columns) using single-molecule RNA *in situ* hybridization (smISH). In the KS fetuses, Sertoli cells (SC, green arrows) and Leydig cells (LC, black arrows) were positive for *XIST* expression, whereas the germ cells (GCs) showed a patchy pattern with some cells being positive (red arrows) and some being negative (blue arrows). The latter was also the case in the control, whereas the somatic cells were negative for *XIST*. Immunohistochemistry using an antibody against MAGE-A4 as a marker of pre-spermatogonia is shown in the left column and tubular borders are marked with black dotted lines. The red rectangle indicates the area magnified to the right. Scalebars represent 100 µm for the left and middle pictures and 20 µm for the right pictures.

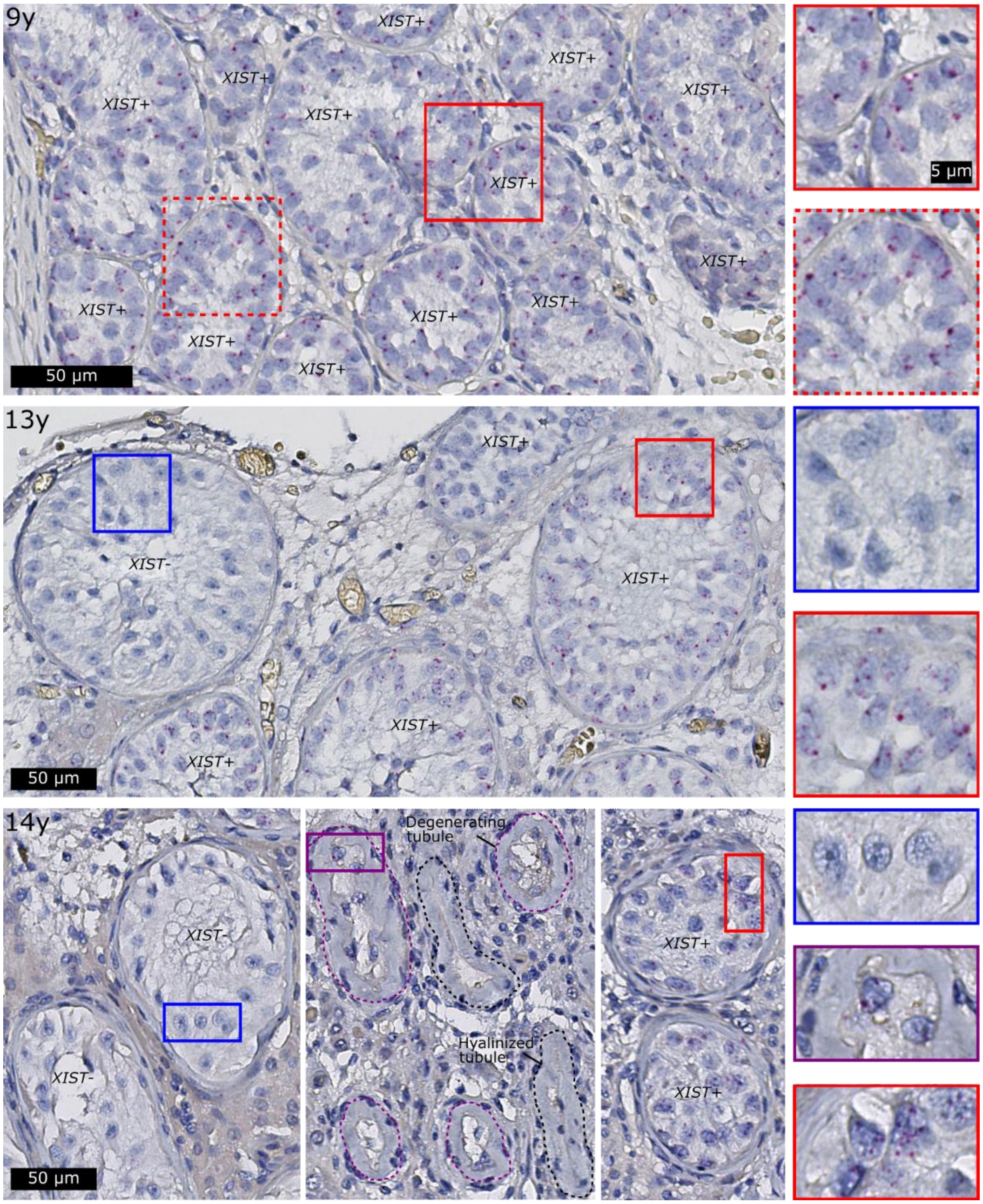


**Fig. S9: *XIST* is expressed in Sertoli cells in prepubertal boys, but at puberty, some Sertoli cells lose *XIST* expression**. RNA single-molecule *in situ* hybridization (smISH) with a probe targeting the *XIST* transcript was performed on biopsies from three boys aged nine, 13 and 14 years old with Klinefelter syndrome (KS). In all samples, Sertoli cells with an immature morphology expressed *XIST* (red squares). In the testis specimens from the 13-year-old and 14-year-old boy, a few tubules, morphologically resembling mature type A showed no expression of *XIST* (blue squares). In the 14-year-old boy, a high proportion of the tubules were hyalinized (black doted lines) or degenerating with a markedly thickened peritubular cell layer and few Sertoli cells left in the tubules (purple dotted lines). These Sertoli cells expressed *XIST* (purple square). Scalebars represent 50 µm 5 µm on the low and high magnifications, respectively.

| Code | Age | Karyotype/diagnosis | Fixative | *XIST*+ SCO tubules | *XIST*- SCO tubules | Tubules with germ cells |
|---|---|---|---|---|---|---|
| aKS1 | 18y | Known KS patient | Stieve | 2 | 7 | 2 |
| aKS2 | 19y | 47,XXY | GR-fix | 4 | 29 | 0 |
| aKS3[*,‖] | 21y | 47,XXY | GR-fix | 29 | 35 | 0 |
| aKS4[*] | 21y | 47,XXY | GR-fix | 21 | 11 | 14 |
| aKS5[†] | 22y | 47,XXY | AFA | 9 | 39 | 0 |
| aKS6 | 24y | 47,XXY | Stieve | 7 | 35 | 11 |
| aKS7[*,†,‡] | 25y | 47,XXY | NBF | 90 | 201 | 16 |
| aKS8[†] | 27y | 47,XXY | AFA | 12 | 5 | 13 |
| aKS9[*] | 27y | 47,XXY | Stieve | 18 | 8 | 0 |
| aKS10 | 27y | 47,XXY | Bouin's | 69 | 8 | 2 |
| aKS11[*,‖] | 28y | 47,XXY | GR-fix | 48 | 165 | 0 |
| aKS12 | 30y | 47,XXY | Stieve | 1 | 7 | 4 |
| aKS13[*] | 30y | Known KS patient | Stieve | 5 | 24 | 0 |
| aKS14[†,§] | 32y | 47,XXY | AFA | 0 | 31 | 0 |
| aNorm1 | 25y | Testicular cancer[¶] | Stieve | 0 | 0 | 30 |
| aNorm2 | 32y | Testicular cancer | GR-fix | 0 | 0 | 29 |
| aNorm3 | 22y | Testicular cancer | Stieve | 0 | 0 | 10 |
| aNorm4[†] | 39y | Testicular cancer | NBF | 0 | 0 | 279 |
| aNorm5[†] | 21y | Testicular cancer | PFA | 0 | 0 | 201 |
| aNorm6[†] | 32y | TESE due to obstructive azoospermia | AFA | 0 | 0 | 132 |
| aCMC | 32y | 46,XY[#] | GR-fix | 0 | 122 | 0 |
| Ovarian tumor | NA | NA | PFA | | | |
| pKS1 | 9y6m | Prepubertal KS patient | Stieve | 508 | 0 | 2 |
| pKS2 | 11y0m | Prepubertal KS patient | Cleland | 342 | 0 | 8 |
| pKS3 | 13y9m | Pubertal KS patient | Stieve | 45 | 4 | 0 |
| pKS4 | 14y6m | Pubertal KS patient | Stieve | 30 | 15 | 0 |
| fKS1 | Gw13 | 47,XXY | NBF | | | |
| fKS2 | Gw14 | 47,XXY | NBF | | | |
| fKS3 | Gw21 | 48,XXYY | NBF | | | |
| fKS4 | Gw22 | 47,XXY | NBF | | | |
| fNorm1 | Gw21 | NA | NBF | | | |
| fNorm2[‖] | Gw20 | NA | NBF | | | |

**Table S1: Sample descriptions.**
*XIST* RNA single-molecule *in situ* hybridization (smISH) was performed on all samples except for aNorm4, aNorm5, and aNorm6 where only DNA smISH was performed. The age refers to the age of the patient at the time of biopsy. The germ cells were either identified by morphological evaluation or MAGE-A4 immunohistochemical staining. KS: Klinefelter syndrome, SCO: Sertoli cell-only, CMC: Cellularity-matched control, Gw: gestational week, NA: not available. [*]Specimens used for pairwise evaluation of type A or B SCO tubules on H&E staining and *XIST* negative or positive tubules on a serial section. [†]Specimens used for DNA chromogen smISH. [‡]Specimen used for DNA XY FISH. [‖]Specimens used for AR, CLU and DEFB119 antibody staining. [§]This specimen only contains *XIST*-negative tubules but was

included in the analysis as its fixative was compatible with DNA chromogen smISH. [¶]Orchiectomy specimens from individuals with testicular cancer. Tissue adjacent to the tumor that was devoid of cancer cells and germ cell neoplasia *in situ* but contained tubules with full spermatogenesis was used. [#]No karyotype was available but there were two gene copies of *SHOX* in the blood of the patient compatible with the karyotype 46,XY.
